# Sensitivity to outcome devaluation in operant tasks is better predicted by food restriction level than reinforcement training schedule in mice

**DOI:** 10.1101/2023.02.23.529699

**Authors:** Maxime Chevée, Courtney J. Kim, Nevin Crow, Emma G. Follman, Erin S. Calipari

**Author notes:** Authors contributed equally to this work (Maxime Chevée, Courtney J. Kim). **Corresponding Author, Erin S. Calipari, PhD**, Associate Director, Vanderbilt Center for Addiction Research, Assistant Professor, Department of Pharmacology, Department of Molecular Physiology and Biophysics, Department of Psychiatry and Behavioral Sciences, Vanderbilt Brain Institute, Vanderbilt University, 865F Light Hall, 2215 Garland Avenue, Nashville, TN 37232.

## Abstract

Behavioral strategies are often classified based on whether reinforcement is controlled by the value of the reinforcer. Value-sensitive behaviors, in which animals update their actions when reinforcer value is changed, are classified as goal-directed; conversely, value-insensitive actions, where behavior remains consistent when the reinforcer is removed or devalued, are considered habitual. Understanding the features of operant training that bias behavioral control toward either strategy is essential to understanding the cognitive and neuronal processes on which they rely. Using basic reinforcement principles, behavior can be biased toward relying on either process: random ratio (RR) schedules are thought to promote the formation of goal-directed behaviors while random intervals (RI) promote habitual control. However, how the schedule-specific features of these task structures relate to external factors to influence behavior is not well understood. Using male and female mice on distinct food restriction levels, we trained each group on RR schedules with responses-per-reinforcer rates matched to their RI counterparts to control for differences in reinforcement rate. We determined that food restriction level has a stronger effect on the behavior of mice following RR schedules than mice following RI schedules and that food restriction better predicted sensitivity to outcome devaluation than training schedule. Our results support the idea the relationships between RR or RI schedules with goal-directed or habitual behaviors, respectively, are more nuanced than previously appreciated and suggest that an animal’s engagement in a task must be accounted for, together with the structure of reinforcement schedules, to appropriately interpret the cognitive underpinnings of behavior.

**Significance statement:** Understanding the basic learning principles that control behavior is essential to developing therapies for psychiatric disorders such as addiction or obsessive-compulsive disorder. Reinforcement schedules are thought to control the reliance on habitual versus goal-directed control during adaptive behaviors. However, external factors that are independent of training schedule also influence behavior, for example by modulating motivation or energy balance. In this study, we find that food restriction levels are at least equally important as reinforcement schedules in shaping adaptive behavior. Our results add to the growing body of work showing the distinction between habitual and goal-directed control is nuanced.

## Introduction

Understanding how animals learn to control their actions is a fundamental goal of behavioral science. One useful framework classifies operant behaviors into two categories: those that are driven by the pursuit of a valuable goal and those that rely on value-insensitive processes (Graybiel, 2008; Smith & Graybiel, 2014). The assessment of these two types of behavior is done in operant conditioning tasks where behavior is maintained by the presentation of an appetitive reinforcer – typically sucrose or food. By devaluating the reinforcer, either by providing *ad libitum* access to it or by pairing it with a negative consequence before the operant session, it is possible to determine whether a behavior is goal-directed: animals whose response rate drops following devaluation are considered to be implementing a goal-directed strategy while those who continue to respond are thought to use a non-goal-directed strategy (often referred to as habitual) (Dickinson et al., 1983; Gremel & Costa, 2013; Killcross & Coutureau, 2003; Schreiner et al., 2020). The cognitive and neuronal processes distinguishing these two types of behaviors are thought to rely on distinct mechanisms (Balleine & Dickinson, 1998; Graybiel & Grafton, 2015; Yin & Knowlton, 2006). Thus, this behavioral framework has guided a large amount of work in behavioral and neuroscience research.

Two types of reinforcement schedules are known to engender different sensitivity to reinforcer devaluation (Dickinson et al., 1983). In random ratio (RR) schedules, a reinforcer is delivered after a certain number of responses; this generates behavior that is goal-directed and sensitive to reinforcer value. Alternatively, random interval (RI) schedules allow access to a reinforcer after a certain amount of time has elapsed. In interval schedules, the delivery of the reinforcer still requires an operant response; however, responses are only reinforced following an elapsed period (Ferster & Skinner, 1957). These schedules often generate behavior over time that is insensitive to changes in the reinforcer. Although this dichotomy is a useful framework to study these types of behaviors, it has been challenged by experimental observations (Schreiner et al., 2020). For example, there is conflicting evidence regarding whether extended training on RR schedules becomes devaluation-insensitive over time (Adams, 1982; Adams & Dickinson, 1981; Dickinson, 1985; Miyachi et al., 1997) or remains goal-directed (Colwill & Rescorla, 1985, 1988; Colwill & Triola, 2002; Garr et al., 2021). Similarly, specific manipulations of the intervals used in RI schedules produce behaviors that remain sensitive to reinforcer devaluation (Garr et al., 2020). However, even when the reinforcement schedules are appropriately tuned, schedule-independent factors – such as food restriction or stress - can also have effects on behavior that could differ depending on the schedule under which the behavior occurs.

It is well-known that external factors can differentially influence behavior that is maintained on different schedules. One prominent example of this is early work showing that the same dose of systemic pentobarbitol can increase or decrease the rate of food reinforcement depending on whether food delivery was reinforced under a ratio or interval schedule, respectively (Dews, 1955, 1956). In the behaviors discussed above – which are maintained by caloric foods - one particularly important factor is the hunger state of the animal. Indeed, food intake is often restricted to increase response rates in operant tasks. While varying food restriction levels affects response rates and response patterns under certain conditions (Blakely & Schlinger, 1988; Lowe et al., 1974; Powell, 1969; Schlinger et al., 1990; Sidman & Stebbins, 1954), the effects of this common manipulation on RR compared to RI schedules are not known. We hypothesized that, because animals following RR schedules control their own rate of reinforcement as opposed to those on RI schedules in which they cannot acquire reinforcers faster than the interval allows, food restriction levels influence performance on RR schedules more strongly than performance on RI schedules and contribute to sensitivity to outcome devaluation independently of schedule.

To explore this idea, we restricted the daily food intake of mice to various degrees and trained them on either RR or RI schedules for which we adjusted the number of responses required in RR schedules to match the actions-per-outcome mice of each restriction group achieved on an RI schedule. This strategy allowed us to mitigate the effect of differences in reinforcement efficiency and to compare the effects of food deprivation levels on performance and value sensitivity across RI and RR schedules. We found that deprivation had a combinatorial effect on operant behavior: increased food restriction was more effective at increasing response rates under RR compared to RI schedules. Furthermore, devaluation tests showed that as animals became more food-deprived, they behaved in a more goal-directed manner, regardless of schedule. Our results add to the growing evidence that the relationship between RR/RI and goal-directed/habitual behavior is far from clear cut and suggest that external factors such as food restriction level must be accounted for, together with the structure of reinforcement schedules, to appropriately interpret the cognitive underpinnings of behavior.

## Materials and methods

### Subjects

Experiments were approved by the Institutional Animal Care and Use Committee of Vanderbilt University Medical Center and conducted according to the National Institutes of Health guidelines for animal care and use. Seventy 8-week-old animals were used for this study. C57BL/6J mice (35 males and 35 females) were acquired from Jackson Laboratory (Bar Harbor, ME; SN: 000,664) and maintained on an 8am/8pm 12-h reverse light cycle. Experiments were performed during the dark phase. Animals were single housed for the duration of the study with unlimited access to water.

The experiments were performed in four sequential training sessions.

**1)** 20 mice (10 females, 10 males) were mildly food restricted (3g/day) for one week. They were then trained on FR1 until acquisition criteria were met (see **Procedure** below) and assigned to a RR10/RR20 schedule or RI30/RI60 schedule such that time to acquisition and sex were balanced across the two groups. This training session resulted in N = 5 females, 5 males performing an RR10/RR20 task under “Mild Restriction” conditions and N = 4 females, 5 males performing an RI30/RI60 task under “Mild Restriction” conditions (one female failed to acquire FR1 for 10 consecutive days).

**2)** 20 mice (10 females, 10 males) were mildly food restricted (3g/day) for one week. They were then trained on FR1 until acquisition criteria were met (see **Procedure** below) and assigned to either a “No Restriction” group, which had unlimited access to food, or to a “Strong Restriction” group which received 2g chow per day, such that time to acquisition and sex were balanced across the two groups. All animals went on to perform an RI30/RI60 task. This training session resulted in N = 5 females, 5 males performing an RI30/RI60 task under “No Restriction” conditions and N = 5 females, 5 males performing an RI30/RI60 task under “Strong Restriction” conditions.

**3)** 20 mice (10 females, 10 males) were mildly food restricted (3g/day) for one week. They were then trained on FR1 until acquisition criteria were met (see **Procedure** below) and assigned to either a “No Restriction” group, which had unlimited access to food, or to a “Strong Restriction” group which received 2g chow per day, such that time to acquisition and sex were balanced across the two groups. Mice assigned to the “No Restriction” group then followed an RR2.8/RR3.4 schedule (ratios matched to the performance of the “No Restriction” group on RI30/RI60) and mice assigned to the “Strong Restriction” group followed an RR5.1/RR9.3 schedule (ratios matched to the performance of the “Strong Restriction” group on RI30/RI60). This training session resulted in N = 4 females, 5 males performing an RR2.8/RR3.4 task under “No Restriction” conditions (one female failed to acquire FR1 for 10 consecutive days) and N = 3 females, 5 males performing an RR5.1/RR9.3 task under “Strong Restriction” conditions (one female was a statistical outlier and one female was erroneously given unlimited access to food in the middle of the experiment and removed from the cohort).

**4)** 10 mice (5 females, 5 males) were mildly food restricted (3g/day) for one week. They were then trained on FR1 until acquisition criteria were met (see **Procedure** below) and then followed a RR3.4/RR5.2 schedule (ratios matched to the performance of the “Mild Restriction” group on RI30/RI60). This training session resulted in N = 5 females, 5 males performing an RR3.4/RR5.2 task under “Mild Restriction” conditions.

### Apparatus

Mice were trained and tested daily in individual standard-wide mouse operant conditioning chambers (Med Associates Inc., St. Albans, Vermont) outfitted with a retractable lever. A 3D-printed divider was inserted in each chamber, limiting the available space to a small square area providing access to a sucrose port and a lever (area: 13 × 13 = 169 cm^2). A custom-made 3D-printed wall insert was used to hold and display a stainless-steel cannula (lick port, 18 gauge, 0.042” ID, 0.05” OD, 0.004”wall thickness), which was connected to a syringe pump for sucrose delivery. An illumination light was affixed above the lick port.

### Procedure

#### General Procedural Information

All sessions lasted until the maximum number of rewards was obtained (51) or 1 hour was reached, whichever came first. White noise signaled the beginning of the session and was on for the entire duration of the session.

#### Task Design

Mice were first trained on a fixed ratio 1 (FR1) schedule of reinforcement. Each lever press resulted in the delivery of 8 μl of a 10% sucrose solution. Additional presses performed after sucrose delivery but before sucrose collection had no programmed consequence and did not count toward obtaining the next reinforcer. Once the reinforcer was collected, lever presses counted again. Mice were considered to have met the acquisition criteria once they had obtained 51 rewards within the allotted 1 h for two consecutive days. Upon meeting the criteria, mice were assigned to different tasks (see *Subjects* above) such that time to acquisition and sex were balanced across groups.

<u>Random Interval (RI) training:</u> RI schedules consisted of three days operating under an RI30 schedule, in which a lever press resulted in sucrose delivery only after an interval of time averaging 30 seconds had passed since consuming the last reinforcer, followed by four days operating under an RI60 schedule. These intervals were determined in keeping with previous studies, which suggested that this training procedure produces habitual actions (Gremel & Costa, 2013; Hilario et al., 2007).
<u>Random Ratio (RR) training:</u> For the cohort reported in Figure 1, the RR schedules consisted of three days operating under an RR10 schedule, in which a lever press resulted in sucrose delivery only after a specific amount of responses (10 on average) had been performed, followed by four days operating under an RR20 schedule. For the three cohorts with distinct food restriction levels, the ratios were calculated to match the responses-per-reinforcer ratio of their counterparts who performed the RI task and are detailed in the Result section (Figure 2B,C) as well as in *Subject* above.

**Figure 1.**
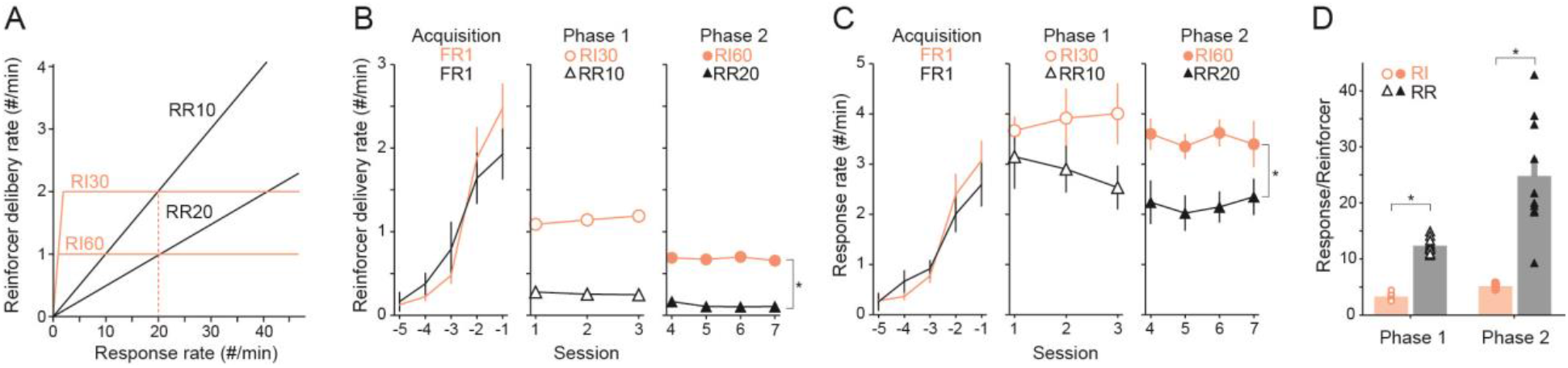
RI30/RI60 and RR10/RR20 produce different operant behavior and different response-per-reinforcer ratios. **(A)** Schematic showing the relationship between response rate and reinforcer delivery rate under 4 example schedules. The red dotted line indicates the mean response rate a subject needs to perform in order to achieve equal reinforcer delivery rates under RI30 or RR10 schedules. The value is the same for achieving equal reinforcer delivery rates between RI60 and RR20. **(B)** Mean reinforcer delivery rates across training, split by schedule. **(C)** Mean response rates across training, split by schedule. **(D)** Mean responses-per-reinforcer ratio during each phase of training. RI N=9 mice, RR=10 mice, * indicates Mann-Whitney U test p<0.05. Data are shown as mean±sem.

**Figure 2.**
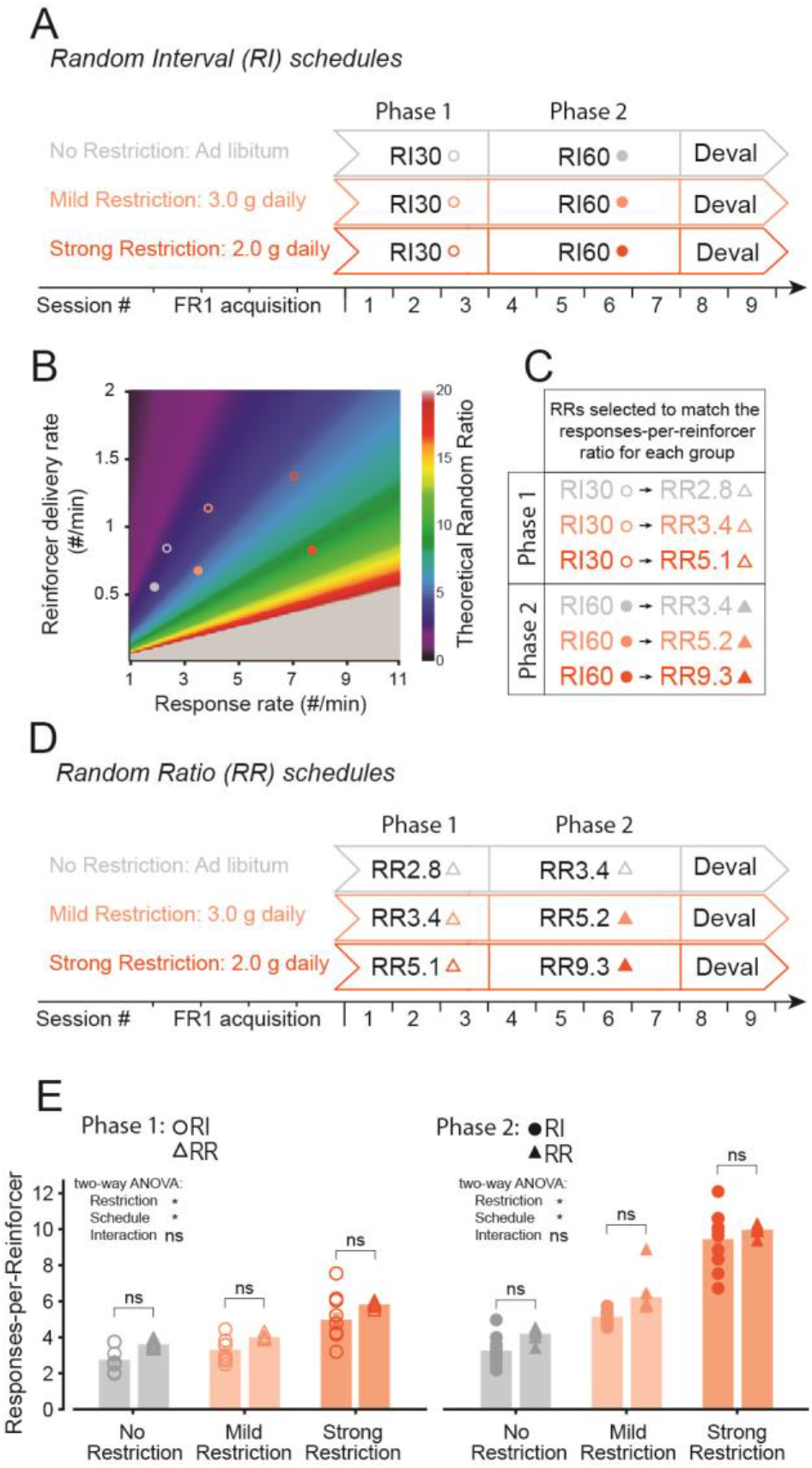
Designing RR schedules to match the responses-per-reinforcer ratios observed in mice following RI schedules under distinct levels of food restriction. **(A)** Schematic showing the training strategy for three groups distinguished by their level of food restriction. Each mouse underwent 3 days of RI30, 4 days of RI60 and 2 days of outcome devaluation testing. **(B)** Plot showing that each pair of response rate/reinforcer delivery rate can be achieved by a specific RR (shown as a color map). The response rate/reinforcer delivery rate values observed for each phase/restriction group are depicted as circles color-coded to match (A). **(C)** Summary of Figure 2B showing the RR that matches each RI based on the performance of each RI group. **(D**) Schematic showing the training strategy for three groups distinguished by their level of food restriction. Each group followed the RR schedule identified in (B-C). **(E)** Mean responses-per-reinforcer ratios for each restriction/schedule group, separated by phase of training. RI-noRestriction N=10 mice, RR-NoRestriction=9 mice, RI-MildRestriction N=9 mice, RR-MildRestriction=10 mice, RI-strongRestriction N=10 mice, RR-strongRestriction=8 mice; * indicates two-way ANOVA p<0.05 (E). ns indicates two-way ANOVA p>0.05 (E) and post hoc Tukey HSD p>0.5 (E). Data are shown as mean±sem.

#### Outcome devaluation tests

Upon completing the seven days of RI or RR training, each mouse underwent an outcome devaluation test. This procedure consists of providing access to the reinforcer (10% sucrose, devalued condition) or to another familiar food source that was not used as a reinforcer (chow, valued condition) *ad libitum* for one hour prior to a 10 minute extinction session, in which the lever presses have no programmed consequences and do not result in reinforcer delivery. Within each cohort, half the mice were assigned to perform this test “Day1: valued, Day2: devalued” or “Day1: devalued, Day2: valued”. Mice which consumed less than 0.5g during either pre-feeding sessions were excluded from the analysis of outcome devaluation tests. A devaluation index was calculated for each mouse (response rate during valued session / (response rate during valued session + response rate during devalued session)), such that an index close to 1 indicates sensitivity to outcome devaluation (ie. goal-directed control).

### Analysis

#### Statistical Analyses

All analyses were performed using custom code in Python (v3.6.13). The SciPy package (v1.5.3) was used to perform paired Mann-Whitney U-tests (Figures 1B-D) and the Pingouin package (v0.3.12) was used to perform two-way ANOVAs as well as the corresponding post-hoc Tukey tests (Figures 2D, 3B, 3D, 4B). The Pearson correlation coefficient (r) and the associated p-value (Figure 4C,D) were calculated using the stats module in SciPy package (v1.5.3). The correlation coefficient r is defined as:

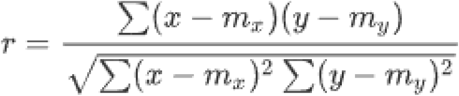

where m_x_ = mean of the vector x, m_y_ = mean of the vector y. The probability density function of the sample correlation coefficient r is defined as (Student 1908):

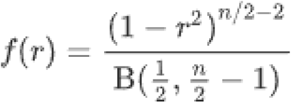

where n = number of samples, B = beta function. The p-value is calculated as the probability that abs(r*) for two random samples drawn from two independent populations is greater or equal to abs(r). All data are reported as Mean ± SEM and all statistical tests used are specified in the Results section.

**Figure 3.**
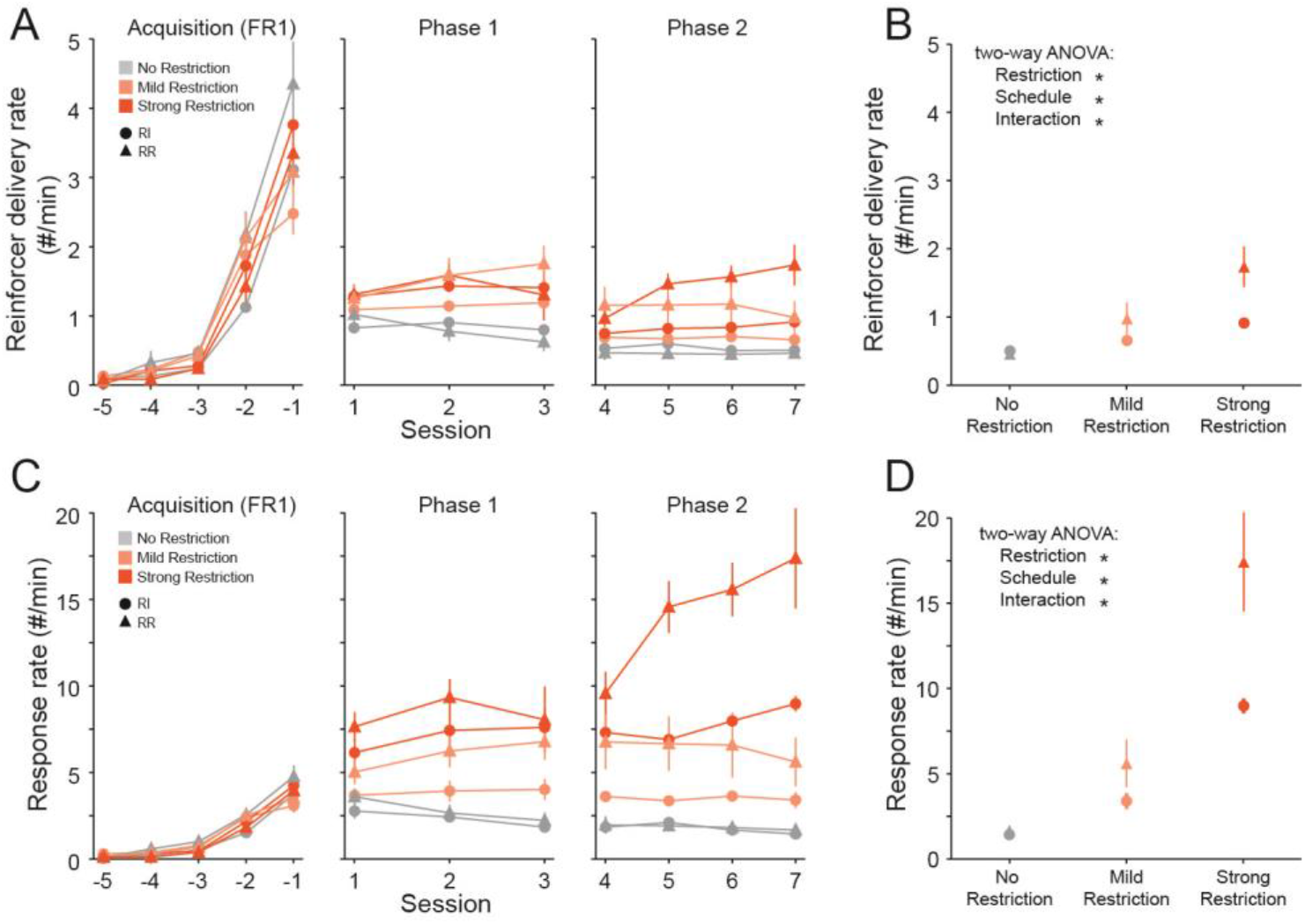
Food restriction increases response rates more effectively in mice following RR schedules than in mice following RI schedules. **(A)** Mean reinforcer delivery rates across training, split by restriction and schedule. **(B)** Mean reinforcer delivery rates on the last day of training summarizing the data used to perform a two-way ANOVA. **(C)** Mean response rates across training, split by restriction and schedule. **(D)** Mean response rates on the last day of training summarizing the data used to perform a two-way ANOVA. RI-noRestriction N=10 mice, RR-NoRestriction=9 mice, RI-MildRestriction N=9 mice, RR-MildRestriction=10 mice, RI-strongRestriction N=10 mice, RR-strongRestriction=8 mice; * indicates two-way ANOVA p<0.05 (B,D). Significant post hoc Tukey HSD test are reported in the result section. Data are shown as mean±sem.

**Figure 4.**
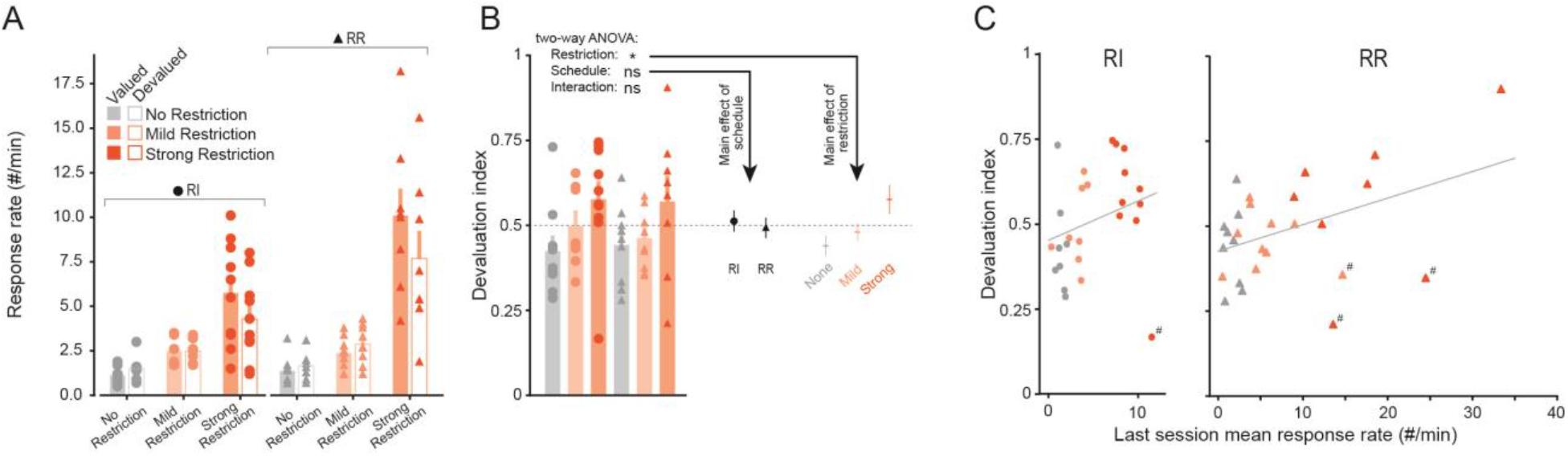
Food restriction is a better predictor of sensitivity to reinforcer devaluation than schedule of reinforcement. **(A)** Summary of response rates during devaluation sessions. Valued conditions consisted of 1 hour access to regular chow and devalued 1 hour access to 10% sucrose. The two test sessions were performed on consecutive days and the order counterbalanced within each group. (valued-RI-NoRestriction: 1.19±0.15 responses/min; devalued-RI-NoRestriction: 1.59±0.23 responses/min; valued-RI-MildRestriction: 2.40±0.26 responses/min; devalued-RI-MildRestriction: 2.44±0.22 responses/min; valued-RI-StrongRestriction: 5.74±0.92 responses/min; devalued-RI-StrongRestriction: 4.28±0.73 responses/min; valued-RR NoRestriction: 1.37±0.27 responses/min; devalued-RR-NoRestriction: 1.67±0.23 responses/min; valued-RR-MildRestriction: 2.51±0.24 responses/min; devalued-RR-MildRestriction: 3.07±0.38 responses/min; valued-RR-StrongRestriction: 10.1±1.53 responses/min; devalued-RR-StrongRestriction: 7.69±1.54 responses/min) **(B)** Same data as shown in (A) summarized as a devaluation index for each mouse (valued response rate / (valued response rate + devalued response rate)). Response rate means for the mice grouped by schedule and by restriction are also shown on the right. **(C)** Plot showing the correlation between the response rate on the last session and devaluation index for each mouse, split by schedule (left: RI, right: RR). The grey line represents a linear regression fit through the data plotted and specific data points discussed on the result section are marked by #. RI-noRestriction N=8 mice, RR-NoRestriction=9 mice, RI-MildRestriction N=8 mice, RR-NoRestriction=10 mice, RI-strongRestriction N=10 mice, RR-strongRestriction=8 mice); * indicates two-way ANOVA p<0.05 (B). ns indicates two-way ANOVA p>0.05 (B). Data are shown as mean±sem.

## Results

### RI and RR schedules in mice can produce different response rates and different responses-per-reinforcer ratios

The stronger relationship between action and outcome in RR schedules compared to RI schedules is often cited as the primary reason for differences in sensitivity to outcome devaluation (Dickinson, 1985; Pérez et al., 2019) (although see (Garr et al., 2020)). However, the two schedules differ in the reinforcer delivery rates they produce at any given response rate, such that mice on RR can increase their reinforcer rate by increasing their response rate while the rate of reinforcer delivery on RI schedules does not correlate with response rate (**Figure 1A**). Therefore, an animal’s response rate can result in distinct responses-per-reinforcer ratios that are dependent on schedule. To illustrate this phenomenon, we compared food-restricted mice trained on RI and RR schedules using commonly used ratios/intervals (Gremel & Costa, 2013; Hilario et al., 2007). First, we restricted the mice’s daily food intake to 3g per day to increase their engagement in the task and trained them to respond for sucrose under a Fixed-Ratio 1 (FR1) schedule of reinforcement. Upon acquisition of FR1, mice went on to a two-phase schedule comprised of either 1.) three days of RI30 followed by four days of RI60 or 2.) three days of RR10 followed by four days of RR20. When reinforcer delivery rate (**Figure 1B**, RI: 0.64±0.06 reinforcers/minute, N=9; RR: 0.10±0.02 reinforcers/minute, N=10) and response rate were assessed (**Figure 1C**, RI: 3.29±0.42 responses/minute, N=9; RR: 2.35±0.36 responses/minute, N=10), we found that by the end of training the RI group had both a faster reinforcer delivery rate (Mann-Whitney U test p=1.9e-4) and a faster response rate (Mann-Whitney U test p=0.033). This result indicates that, under food restriction conditions, an RI30/RI60 training schedule is more effective than a RR10/RR20 schedule at increasing reinforcer and response rates.

Interestingly, the response-per-reinforcer ratio was much larger for the RR group than for the RI group during both phases of training (**Figure 1D**, Phase 1 - RI: 3.33±0.21, N=9; RR: 12.3±0.45, N=10, Mann-Whitney U test p=1.4e-4; Phase 2 - RI: 5.11±0.12, N=9; RR: 24.8±3.19, N=10, Mann-Whitney U test p=1.4e-4). In other words, reinforcer rates between RR and RI schedules were not comparable (they would only have been comparable at 20 responses/min, **Figure 1A** - dotted line). Thus, the differences in behavior following training under RI vs RR tasks are confounded with differences in response rates, which themselves are determined by reinforcement rate. We next set out to test whether the reinforcement rate or the schedule contingency better predicted operant performance and value sensitivity.

### Designing RR schedules to match the responses-per-reinforcer ratios observed in mice following RI30/RI60 schedules under distinct levels of food restriction

To determine whether differences in response rates rather than differences in schedule contingencies best explain sensitivity to reinforcer devaluation, we first had to tailor RR schedules to produce a similar relationship between response rate and reinforcer delivery (responses-per-reinforcer) compared to RI schedules. To this end, three groups were trained on RI30/RI60 schedules under different food restriction conditions (No Restriction, Mild Restriction, Strong Restriction, **Figure 2A**). Animals had either unlimited access to food in their home cage (no restriction), received 3g of chow per day (Mild restriction) or received only 2g of chow per day (Strong restriction). For each group and each phase of training, we determined the mean response rate as well as the mean reinforcer delivery rate and computed the RR schedule at which the responses-per-reinforcer ratio would match that of the RI schedule given the observed response rates (defined as response rate / reinforcer delivery rate - **Figure 2B,C**).

We then trained three groups on the identified RI-matched RR schedules for each food restriction condition (**Figure 2D**) and compared the responses-per-reinforcer ratios during each phase of training to validate our strategy. During phase 1, a two-way ANOVA revealed that there was a main effect of both the restriction group and of task schedule (RI_NoR: 2.74±0.19, N=10; RR_NoR: 3.61±0.06, N=9; RI_MildR: 3.33±0.21, N=9; RR_MildR: 4.00±0.04, N=10; RI_StrongR: 4.98±0.40, N=10; RR_StrongR: 5.80±0.05, N=8; two-way ANOVA, effect of task: F=19.8, p=4.8e-05; effect of restriction: F=57.1, p= 1.25e-13), however posthoc comparisons showed there were no significant differences between each pair of restriction-matched RI-RR group (RI_NoR-versus-RR_NoR Tukey HSD p-adjuted=0.065; RI_MildR-versus-RR_mildR Tukey HSD p-adjuted=0.25; RI_StrongR-versus-RR_StrongR Tukey HSD p-adjuted=0.11). Results during phase two were similar (RI_NoR: 3.25±0.25, N=10; RR_NoR: 4.18±0.11, N=9; RI_MildR: 5.11±0.12, N=9; RR_MildR: 6.17±0.31, N=10; RI_StrongR: 9.44±0.51, N=10; RR_StrongR: 9.97±0.1, N=8; two-way ANOVA, effect of task: F=12.5, p=8.88e-04; effect of restriction: F=168, p=2.54e-25) and posthoc tests again revealed no differences between each pair of restriction-matched RI-RR group (RI_NoR-versus-RR_NoR Tukey HSD p-adjuted=0.23; RI_MildR-versus-RR_mildR Tukey HSD p-adjuted=0.12; RI_StrongR-versus-RR_StrongR Tukey HSD p-adjuted=0.78). Thus, our training strategy successfully produced three pairs of RI-RR groups with distinct levels of food restrictions and matched responses-per-reinforcer ratios, allowing us to compare the effects of restriction and task schedule on behavior and outcome devaluation.

### Food restriction increases response rates more effectively in mice following RR schedules than in mice following RI schedules

We first analyzed differences in reinforcer delivery rate (**Figure 3A-B**) and response rate (**Figure 3C-D**). We tested how food restriction and task schedule influenced reinforcer delivery by the last day of training (**Figure 3B**, RI_NoR: 0.50±0.05 reinforcers/minute, N=10; RR_NoR: 0.46±0.07 reinforcers/minute, N=9; RI_MildR: 0.64±0.06 reinforcers/minute, N=9; RR_MildR: 0.96±0.21 reinforcers/minute, N=10; RI_StrongR: 0.91±0.02 reinforcers/minute, N=10; RR_StrongR: 1.74±0.3 reinforcers/minute, N=8) and found that both restriction and task schedule had a main effect on final reinforcer delivery rate (two-way ANOVA, effect of task: F=8.97, p=4.3e-03; effect of restriction: F=15.4 p=6.0e-6). In addition, the interaction term was significant (two-way ANOVA, interaction between task and restriction F=4.40, p=0.017), and post hoc tests revealed that only the Strong Restriction group performing an RR task achieved a higher rate of reinforcer delivery than any other group (RI_NoR-versus-RR_StrongR Tukey HSD p-adjuted=0.0010; RI_MildR-versus-RR_StrongR Tukey HSD p-adjuted=0.0010; RR_NoR-versus-RR_StrongR Tukey HSD p-adjuted=0.0010; RI_StrongR-versus-RR_StrongR Tukey HSD p-adjuted=0.0036; RR_MildR-versus-RR_StrongR Tukey HSD p-adjuted=0.0075). When we tested for effects on response rates, we found similar results (**Figure 3D**, RI_NoR: 1.43±0.19 responses/minute, N=10; RR_NoR: 1.66±0.29 responses/minute, N=9; RI_MildR: 3.29±0.42 responses/minute, N=9; RR_MildR: 5.51±1.25 responses/minute, N=10; RI_StrongR: 8.96±0.44 responses/minute, N=10; RR_StrongR: 17.4±2.90 responses/minute, N=8). The main effects of restriction and task schedule were again significant (two-way ANOVA, effect of task: F=13.3, p=6.4e-04; effect of restriction: F=48.6, p=1.9e-12), as was the interaction term (two-way ANOVA, interaction between task and restriction F=6.37, p=3.4e-3). Similar to the reinforcer delivery rate, post hoc tests showed that the Strong Restriction group performing an RR task had a larger response rate than all other groups (RI_NoR-versus-RR_StrongR Tukey HSD p-adjuted=0.0010; RI_MildR-versus-RR_StrongR Tukey HSD p-adjuted=0.0010; RR_NoR-versus-RR_StrongR Tukey HSD p-adjuted=0.0010; RI_StrongR-versus-RR_StrongR Tukey HSD p-adjuted=0.0010; RR_MildR-versus-RR_StrongR Tukey HSD p-adjuted=0.0010), however here the Strong Restriction group performing an RI task also achieved higher response rates than several groups (RI_NoR-versus-RI_StrongR Tukey HSD p-adjuted=0.0010; RR_NoR-versus-RI_StrongR Tukey HSD p-adjuted=0.0010; RI_MildR-versus-RI_StrongR Tukey HSD p-adjuted=0.015).

We can draw three conclusions from these results. First, they show that when the responses-per-reinforcer ratios are matched, mice trained on RI schedules do not achieve a higher response rate than their counterparts trained on RR schedules under any circumstances. Second, both food restriction and task schedule independently influenced reinforcer delivery rate and response rate, with more restricted mice and mice on RR schedules achieving higher rates than less restricted mice and mice on RI schedules, respectively. Third, there was a clear combinatorial effect of restriction and task schedule, such that food restriction increases reinforcer delivery rates and response rates more effectively in mice following RR schedules than in mice following RI schedules. These results are consistent with the idea that RR schedules generate behaviors that are more sensitive to value than RI schedules.

### Food restriction is a better predictor of sensitivity to reinforcer devaluation than schedule of reinforcement

After completing the seven days of RI or RR training, all mice underwent outcome devaluation sessions (see methods). The raw results of each session for each mouse are depicted in **Figure 4A** and a two-way ANOVA was performed on the devaluation index derived from these data (**Figure 4B** left, RI_NoR: 0.43±0.05, N=8; RR_NoR: 0.44±0.04, N=9; RI_MildR: 0.49±0.04, N=8; RR_MildR: 0.46±0.03, N=10; RI_StrongR: 0.58±0.05, N=10; RR_StrongR: 0.57±0.08, N=8) to identify the effects of task schedule and of food restriction on outcome devaluation. We found that only food restriction had a significant main effect on the devaluation index (two-way ANOVA F=4.06, p=0.024) while task schedule had no effect (two way ANOVA F=0.0678, p=0.80). To illustrate this result, we plotted the means for groups collapsed by restriction or by task schedule (**Figure 4B** right). Taken together, these results show that when the response rates and reinforcer deliveries are matched, restriction level, but not schedule contingency, determines value sensitivity.

Our data thus suggest that the level of food restriction – which controls response rates – is more predictive of reinforcer devaluation than the reinforcement schedule itself. This result predicts that mice with the highest response rates should also have the highest devaluation indices. When we tested this prediction, we found that the response rate on the last training session and devaluation index were positively correlated (**Figure 4C** left, Pearson r=0.27, slope of fit=0.012), although not significantly (p=0.18) for mice trained on RI schedules and that they were also positively correlated for mice trained on RR schedules (**Figure 4C** right, Pearson r=0.43, slope of fit=0.0079, p=0.024). It is noteworthy that all mice that degraded the strength of the correlation (marked by # in **Figure 4C**) underwent the sucrose devaluation extinction session (devalued condition) before the food devaluation session (valued condition). Because these animals extinguished, their devalued response rate was high and their valued response rate was low, leading to a devaluation index artificially lowered by the order in which the test was performed. When these mice were excluded from the analysis presented in **Figure 4C**, the correlation between the response rate on the last session and devaluation index was positive and strong for both mice trained on RI schedules (Pearson r=0.54, slope of fit=0.022, p=0.0057) and mice trained on RR schedules (Pearson r=0.78, slope of fit=0.014, p=8.2e-6). Thus, the more a mouse responds, the more likely it is to appear sensitive to reinforcer devaluation.

## Discussion

Our results show that the level of food restrictions under which animals learn and operate affects their performance differently based on the type of reinforcement schedule they are following. Indeed, food restriction increased response rates and reinforcer delivery rates more strongly in mice following RR schedules compared to mice following RI schedules. In parallel, animals were rendered more value-driven by increasing levels of food restriction. Our study therefore reveals that the relationship between schedule, value and behavior is complex. The task schedule fundamentally interacted with food restriction level to shape behavioral output, yet it showed little effect on value-sensitivity compared to food restriction level. Thus, schedule of reinforcement must be considered in combination with other factors that may influence motivation when attempting to identify the cognitive basis underlying operant behaviors.

Our results are consistent with previous studies in rats (Dawson & Dickinson, 1990), pigeons (Catania et al., 1977; Peele et al., 1984; Zuriff, 1970) and humans (McDowell & Wixted, 1986; Pérez et al., 2019; Wit et al., 2018) showing that when reinforcer delivery rate is matched, animals following RR schedules generally achieve higher response rates than those following RI schedules. Both the preferential reinforcement of long inter-response times (IRT) in RI schedules (Krame & Rilling, 1970; Kuch & Platt, 1976) and the stronger action-outcome correlation in RR schedules (Dickinson, 1985; Ferster & Skinner, 1957; Pérez et al., 2019 although see Garr et al., 2020) are thought to contribute to generating this difference, and our finding that response rates are more sensitive to food restriction level in mice trained on RR schedules compared to RI schedules complements these interpretations. Furthermore, previous work has shown that while training schedule has little effects on motivation (Johnson et al., 2022), food restriction levels do impact it (Dietze et al., 2016; Malikovic et al., 2018). By manipulating food restriction levels, our findings provide further support to the notion that the ability to control one’s own reinforcement rate explains the difference in response rates generated by RR and RI schedules.

Several studies have explored the effects of changes in reinforcer value or in food restriction levels on performance (Balleine, 1992; Blakely & Schlinger, 1988; Lowe et al., 1974; Powell, 1969; Schlinger et al., 1990; Sidman & Stebbins, 1954). However, these studies were largely focused on evaluating the effects of acute changes in value rather than understanding the systematic relationship between food restriction and schedule. Nevertheless, changes in reinforcer value have been shown to have a stronger effect on actions temporally close to the reinforcer delivery compared to those further away (Dickinson & Balleine, 1995; Holland & Rescorla, 1975). Our findings support this idea because they suggest that tasks that promote a high response rate, either via food restriction or via denser schedules, are more likely to appear sensitive to value. This is also supported by the observation that leaner RI schedules are more likely to produce devaluation-insensitive behaviors (Garr et al., 2020) and that as long as animals continue to perform and remain in control of their own reinforcement rate, they remain goal-directed even for long periods of time (Colwill & Rescorla, 1985, 1988; Colwill & Triola, 2002; Corbit et al., 2012; Garr et al., 2021). Overall, our findings support the idea that schedule-dependent and -independent factors interact to influence behavior.

Finally, our results add to the growing evidence against a clear-cut relationship between schedule and cognitive basis of behavior, in which RR training produces goal-directed actions and RI training produces habitual actions (Colwill & Rescorla, 1990; Dezfouli & Balleine, 2012; Garr & Delamater, 2019; Schreiner et al., 2020; Wit et al., 2018). While the two schedules produce inherently distinct features such as reinforcement uncertainty (de Russo et al., 2010), the temporal distribution of reinforcers and of responses (Adams, 1982; Doughty & Lattal, 2003; Urcelay & Jonkman, 2019), the correlation between action and outcome or the preferential reinforcement of long IRTs (Krame & Rilling, 1970; Kuch & Platt, 1976), that all likely contribute to differences in value sensitivity, external factors are at least equally important. Our data show that food restriction is in fact a better predictor of sensitivity to outcome devaluation than training schedule and support evidence that external factors such as reinforcer value, on-board drug or stimulus saliency (Vandaele et al., 2017), both contribute to, and can fundamentally control, the underlying cognitive basis of behavior.

## Acknowledgement

This work was supported by NIH grants DA042111 and DA048931 to E.S.C., 5T32MH065215-18 to M.C, as well as by funds from Brain and Behavior Research Foundation, the Whitehall Foundation, and the Edward Mallinckrodt, Jr. Foundation to E.S.C.

## References

Adams, C. D. (1982). Variations in the sensitivity of instrumental responding to reinforcer devaluation. The Quarterly Journal of Experimental Psychology Section B, 34(2), 77–98.

Adams, C. D., & Dickinson, A. (1981). Instrumental Responding Following Reinforcer devaluation. Quarterly Journal of Experimental Psychology, 33, 109–121.

Balleine, B. (1992). Instrumental Performance Following a Shift in Primary Motivation Depends on Incentive Learning. Journal of Experimental Psychology: Animal Behavior Processes, 18(3), 236–250.

Balleine, B., & Dickinson, A. (1998). Goal-directed instrumental action: contingency and incentive learning and their cortical substrates. Neuropharmacology, 37, 407–419.

Blakely, E., & Schlinger, H. (1988). Determinants of pausing under variable-ratio schedules: reinforcer magnitude, ratio size, and schedule configuration. Journal of the Experimental Analysis of Behavior, 1(1), 65–73.

Catania, A. C., Matrrhews, T. J., Silverman, P. J., & Yohalem, R. (1977). Variable-interval responding. Journal of Experimental Analysis of Behavior, 2, 155–161.

Colwill, R. M., & Rescorla, R. A. (1985). Instrumental responding remains sensitive to reinforcer devaluation after extensive training. Journal of Experimental Psychology: Animal Behavior Processes, 11(4), 520–536.

Colwill, R. M., & Rescorla, R. A. (1988). The role of response-reinforcer associations increases throughout extended instrumental training. Animal Learning & Behavior, 16(1), 105–111.

Colwill, R. M., & Rescorla, R. A. (1990). Evidence for the hierarchical structure of instrumental learning. Animal Learning & Behavior, 18(1), 71–82.

Colwill, R. M., & Triola, S. M. (2002). Instrumental responding remains under the control of the consequent outcome after extended training. Behavioural Processes, 57(1), 51–64.

Corbit, L. H., Nie, H., & Janak, P. H. (2012). Habitual alcohol seeking: Time course and the contribution of subregions of the dorsal striatum. Biological Psychiatry, 72(5), 389–395.

Dawson, G. R., & Dickinson, A. (1990). Performance on Ratio and Interval Schedules with Matched Reinforcement Rates. The Quarterly Journal of Experimental Psychology Section B, 42(3), 225–239.

de Russo, A. L., Fan, D., Gupta, J., Shelest, O., Costa, R. M., & Yin, H. H. (2010). Instrumental uncertainty as a determinant of behavior under interval schedules of reinforcement. Frontiers in Integrative Neuroscience, 4(May), 1–8.

Dews, P. B. (1955). Studies on behavior. I. Differential sensitivity to pentobarbital of pecking performance in pigeons depending on the schedule of reward. Journal of Pharmacology and Experimental Therapeutics, 113(4), 393–401.

Dews, P. B. (1956). Modification by drugs of performance on simple schedules of positive reinforcement. Annals New York Academy of Sciences, 65(4), 268–281.

Dezfouli, A., & Balleine, B. (2012). Habits, action sequences and reinforcement learning. European Journal of Neuroscience, 35(7), 1036–1051.

Dickinson, A. (1985). Actions and habits: the developm ent of behavioural autonom y. Philosophical Transactions of the Royal Society of London. Series B, 308, 67–78.

Dickinson, A., & Balleine, B. (1995). Motivational Control of Instrumental Action. Current Directions in Psychological Science, 4(5), 162–167.

Dickinson, A., Nicholas, D. J., & Adams, C. D. (1983). The effect of the instrumental training contingency on susceptibility to reinforcer devaluation. The Quarterly Journal of Experimental Psychology Section B, 35(1), 35–51.

Dietze, S., Lees, K. R., Fink, H., Brosda, J., & Voigt, J. P. (2016). Food deprivation, body weight loss and anxiety-related behavior in rats. Animals, 6(1).

Doughty, A. H., & Lattal, K. A. (2003). Response persistence under variable-time schedules following immediate and unsignalled delayed reinforcement. Quarterly Journal of Experimental Psychology Section B: Comparative and Physiological Psychology, 56 B(3), 267–277.

Ferster, C. B., & Skinner, B. F. (1957). Schedules of reinforcement. Appleton-Century-Crofts, 79(1911), 5326.

Garr, E., Bushra, B., Tu, N., Delamater, A. R., Garr, E., Bushra, B., Tu, N., & Delamater, A. R. (2020). Goal-Directed Control on Interval Schedules Does Not Depend on the Action – Outcome Correlation. Journal of Experimental Physiology, 46(1), 47–64.

Garr, E., & Delamater, A. R. (2019). Exploring the relationship between actions, habits, and automaticity in an action sequence task. Learning and Memory, 26(4), 128–132.

Garr, E., Padovan-Hernandez, Y., Janak, P. H., & Delamater, A. R. (2021). Maintained goal-directed control with overtraining on ratio schedules. Learning & Memory (Cold Spring Harbor, N.Y.), 28(12), 435–439.

Graybiel, A. M. (2008). Habits, rituals, and the evaluative brain. Annual Review of Neuroscience, 31, 359–387.

Graybiel, A. M., & Grafton, S. T. (2015). The striatum: where skills and habits meet. Cold Spring Harbor Perspectives in Biology, 7(8), a021691.

Gremel, C. M., & Costa, R. M. (2013). Orbitofrontal and striatal circuits dynamically encode the shift between goal-directed and habitual actions. Nature Communications, 4, 2264.

Hilario, M. R. F., Clouse, E., Yin, H. H., & Costa, R. M. (2007). Endocannabinoid signaling is critical for habit formation. Frontiers in Integrative Neuroscience, 1(November), 1–12.

Holland, P. C., & Rescorla, R. A. (1975). The Effect of Two Ways of Devaluing the Unconditioned Stimulus after First-and Second-Order Appetitive Conditioning. Journal of Experimental Psychology: Animal Behavior Processes, 1(4), 355–363.

Johnson, A. R., Christensen, B. A., Kelly, S. J., & Calipari, E. S. (2022). The influence of reinforcement schedule on experience-dependent changes in motivation. 1–11.

Killcross, S., & Coutureau, E. (2003). Coordination of Actions and Habits in the Medial Prefrontal Cortex of Rats. Cerebral Cortex, 2, 400–408.

Krame, T. J., & Rilling, M. (1970). Differential reinforcement of low rates: a selective critique. Psychological Bulletin, 74(4).

Kuch, D., & Platt, J. R. (1976). Reinforcement rate and interresponse time differentiation. Journal of the Experimental Analysis of Behavior, 3(3), 471–486.

Lowe, C. F., Davey, G. C. L., & Harzem, P. (1974). Effects of reinforcement magnitude on interval and ratio schedules. Journal of the Experimental Analysis of Behavior, 3(3), 553–560.

Malikovic, J., Feyissa, D. D., Hussein, A. M., Höger, H., Lubec, G., & Korz, V. (2018). Moderate differences in feeding diets largely affect motivation and spatial cognition in adult and aged but less in young male rats. Frontiers in Aging Neuroscience, 10(AUG), 1–8.

McDowell, J. J., & Wixted, J. T. (1986). Variable-ratio schedules as variable-interval schedules with linear feedback loops. Journal of the Experimental Analysis of Behavior, 3.

Miyachi, S., Hikosaka, O., Miyashita, K., Karadi, Z., & Rand, M. K. (1997). Differential roles of monkey striatum in learning of sequential hand movement. Experimental Brain Research, 115, 1–5.

Peele, D. B., Casey, J., & Silberberg, A. (1984). Primacy of Interresponse-Tirne Reinforcement in Accounting for Rate Differences Under Variable-Ratio and Variable-Interval Schedules. Journal of Experimental Psychology: Animal Behavior Processes, 10(2), 149–167.

Pérez, O. D., Aitken, M. R. F., Zhukovsky, P., Soto, F. A., Urcelay, G. P., & Dickinson, A. (2019). Human instrumental performance in ratio and interval contingencies: A challenge for associative theory. Quarterly Journal of Experimental Psychology, 72(2), 311–321.

Powell, R. W. (1969). The effect of reinforcement magnitude upon responding under fixed-ratio schedules. Journal of the Experimental Analysis of Behavior, 12(4), 605–608.

Schlinger, H., Blakely, E., & Kaczor, T. (1990). Pausing under variable-ratio schedules: interaction of reinforcer magnitude, variable-ratio size, and lowest ratio. Journal of the Experimental Analysis of Behavior, 1(1), 133–139.

Schreiner, D. C., Renteria, R., & Gremel, C. M. (2020). Fractionating the all-or-nothing definition of goal-directed and habitual decision-making. Journal of Neuroscience Research, 98(6), 998–1006. https://doi.org/10.1002/jnr.24545

Sidman, M., & Stebbins, W. C. (1954). Satiation effects under fixed-ratio schedules of reinforcement. Journal of Comparative and Physiological Psychology, 47(2), 114–116.

Smith, K. S., & Graybiel, A. M. (2014). Investigating habits: Strategies, technologies and models. Frontiers in Behavioral Neuroscience, 8(FEB), 1–17.

Student, “Probable error of a correlation coefficient”, Biometrika, Volume 6, Issue 2-3, 1 September 1908, pp. 302–310.

Urcelay, G. P., & Jonkman, S. (2019). Delayed Rewards Facilitate Habit Formation. Journal of Experimental Psychology: Animal Learning and Cognition, 45(4), 413–421.

Vandaele, Y., Pribut, H. J., & Janak, P. H. (2017). Lever insertion as a salient stimulus promoting insensitivity to outcome devaluation. Frontiers in Integrative Neuroscience, 11(September), 1–13.

Wit, S. de, Kindt, M., Knot, S. L., Verhoeven, A. A. C., Robbins, T. W., Gasull-camos, J., & Gillan, C. M. (2018). Shifting the Balance Between Goals and Habits: Five Failures in Experimental Habit Induction. Journal of Experimental Psychology: General, 147(7), 1043–1065.

Yin, H. H., & Knowlton, B. J. (2006). The role of the basal ganglia in habit formation. Nature Reviews Neuroscience, 7(6), 464–476.

Zuriff, G. E. (1970). A comparison of variable-ratio and variable-interval schedules of reinforcement. Journal of Experimental Analysis of Behavior, 3(3), 369–374.

